# A forward-engineered muscle tissue driven soft robotic swimmer

**DOI:** 10.1101/2024.10.30.621139

**Authors:** W. C. Drennan, O. Aydin, B. Emon, Z. Li, M. S. H. Joy, A. Barishman, Y. Kim, M. Wei, D. Denham, A. Carrillo, M. T. A. Saif

**Affiliations:** Department of Mechanical Science and Engineering, University of Illinois at Urbana-Champaign; Urbana, USA; CZ Biohub Chicago, LLC; Chicago, USA; Carl R. Woese Institute for Genomic Biology, University of Illinois at Urbana-Champaign; Urbana, USA; Cullen College of Engineering, Biomedical Engineering, University of Houston; Houston, USA; Department of Bioengineering, University of Illinois at Urbana-Champaign; Urbana, USA; Department of Molecular and Cellular Biology, University of Illinois Urbana-Champaign; Urbana, USA

## Abstract

The integration of biological actuators with soft scaffolds has led to biohybrid robots including microscale flagellate-like swimmers which generate thrust by waving their flagella-like tails. However, they achieve swimming speeds of only 0.014 body lengths per minute, Reynolds number (*Re*) ∼ 10⁻³, which is much slower than natural flagellates (*O*(10^2^ – 10^3^) body lengths per minute). To investigate this, we applied theoretical and experimental methods, including fabrication of a swimmer that converts muscle contractions into large angular tail displacements, reaching swimming speeds of 86.8 μm/s (0.58 body lengths per minute), surpassing low-*Re* predictions. Swimming dynamics sharply transition from a low-Re (∼ 10⁻³) to an intermediate-*Re* (∼ 0.1) regime when the actuation angle exceeded 4°. We used the swimmer to study the ability of muscle to adapt to mechanical stiffness and the beneficial effects of neuromuscular coculture on muscle development. These insights into mechanical and chemical cues will help optimize future biobots.

**Teaser:** How are biological flagellate swimmers like *E. coli* and sperm cells so fast? We have built a new biohybrid robot to explore the theory.

## INTRODUCTION

Biohybrid robots have emerged over the past two decades serving as platforms to study the biomechanics of living actuators as well as novel fabrication and control techniques (*1–5*). Several recently developed muscle-actuated robots, ranging in size scale from 10^-4^ to 10^-2^ meters, have allowed for the basic study of locomotion and emergent behaviors (*6–8*). They include biomimetic swimmers resembling jellyfish (*9*, *10*), manta rays (*11*), and flagellates (*12*, *13*), crawling robots that use structural asymmetry to achieve unidirectional motion (*14–17*), a valveless pumping robot (*18*), and a variety of micromanipulators which function both in (*19*) and out (*20*, *21*) of media. They usually consist of cardiac or skeletal muscle interfaced with a soft scaffold (silicone elastomers, hydrogels) and either operate autonomously or receive pacing through electrical or optogenetic stimulation.

Recent studies have focused on the emergence of living tissues in biobots through the introduction of multiple cell types (*3*, *22*), including motor neurons which can interface with skeletal muscle (*12*, *23–26*). A recently developed flagellate-like biohybrid swimmer demonstrated how these cell types can be combined to form a function motor-unit actuator. The skeletal muscle tissue, coupled to the dynamics of on-board motor neurons, was used to actuate the base of a pair of thin tails. Actuation of the tails elicits a propagating wave. The time-asymmetric nature of these waves allows for net thrust at Reynolds number (*Re*) on the order of 10^-3^. This biobot swam at 0.7 µm/s or 0.014 body lengths per minute, much slower than natural flagellar swimmers, including sperm which swim on the order of 100 body lengths per minute. This motivated us to explore the design space and the associated mechanisms in small-scale biohybrid flagellate-like swimmers that would allow for increased speed. We employ theory, simulation, and experimentation in this work. Note that large scale biohybrid swimmers including a spring-shaped swimmer (*8*) and a cardiac muscle-powered biohybrid stingray (*11*), have been shown to reach relative average velocities of 2.77 and 6.42 body lengths per minute, where the mechanism of thrust generation is different from flagellum-mediated swimming.

Flagellate swimmers operate across a range of flow regimes, including some types of bacteria and algae operating at *Re* ∼ 10^-6^ and sperm operating in the range of *Re ∼* 10^-5^ – 10^-2^ (*27–31*). These biological swimmers generate thrust by moving their flagella in a planar rowing or three-dimensional corkscrew motion, thereby creating time-asymmetric waves that mediate net propulsion at low *Re*. We are interested in producing a biobot propelled by generating planar waves in a thin elastic tail actuated by periodic, linear and angular displacements at its base. The resultant thrust from these waves can be predicted by a low-*Re*, linearized model of slender body elasto-hydrodynamics (*12*, *32*). Our simulations, using this theory, predict a steep increase in thrust from a boundary condition of large angular displacements at the base of the tail, as compared with increasing translation (*33*). Intuitively, improved speed with increased angular actuation follows from a scaling argument about the bending waves. The amplitude of the bending waves is proportional to the amplitude of the linear displacement of the base of the tail and to the product of the length of the tail and the amplitude of angular displacement. For a muscle tissue that can only contract a fixed percentage of its length, changes to the muscle and swimmer size result in increasing drag which cancels out the benefit from increased translation amplitude. It is possible to introduce a compliant rocking mechanism to translate small amplitude muscle contractions into large amplitude angular displacement of the base of the tail at the expense of a larger force resisting the contraction of the muscle. By doing this, it is possible to increase the amplitude of the bending wave proportional to the length of the tails without changing the size of the muscle.

To take full advantage of an appropriate compliant rocking mechanism, it is also necessary to ensure strong maturation of the muscle tissue. The process by which muscle fibers mature, termed myofibrillogenesis, depends on a number of mechanical and chemical factors (*34*). *Mechanical factors*: During the maturation process, myoblast muscle precursor cells fuse into multinucleated nascent myotubes which stretch out and pull on the surrounding matrix. Based on in vivo study of drosophila abdominal muscles which resemble the cross-striated skeletal muscle of mammals, this pulling generates tension and is necessary for the formation of immature contractile units of actin and myosin (*35*). These immature units undergo spontaneous, calcium dependent twitching, which appears to be necessary for the organization of highly periodic cross-striated sarcomeres, the key force generating units in skeletal muscle. This implies that the mechanical stiffness of the mechanism that serves as an anchor for the muscle needs to be sufficient to allow the muscle tension to develop, and yet compliant enough to allow the muscle to twitch. *Chemical factors*: In vitro studies with the immortalized mouse myoblasts cell line, C2C12, demonstrate that this maturation process requires a reduction in growth factors, which can be achieved by reducing the amount of serum in the media (*36*, *37*). A remarkable finding from in vitro studies of neuromuscular junction (NMJ) formation is that the spontaneous twitching of C2C12 derived skeletal muscle increases in the presence of motor neurons (*23*, *38*). We therefore expect that neurons in the vicinity of a muscle culture will improve muscle maturation as demonstrated by an increase in overall contractility and force, although the underlying mechanism of their cross talk remains unclear as well as the culture conditions necessary to allow for it. We consider both mechanical and chemical factors in the development of our swimming biobot.

We present a biohybrid flagellar swimmer that maximizes angular displacement of the base of the tails and improves muscle maturation by offering favorable anchor stiffness and proximity of motor neurons. To our surprise, we find (1) that the speed of the swimmer far exceeds the average velocity (by an order of magnitude) that our model predicts, (2) that the speed shows a transition from low to high values when the tail angle exceeds a threshold value, suggestive of a phase transition, and (3) that muscle maturation is invariant to the history of anchor stiffness such that there seems to be a homeostatic state of the muscle for a given anchor stiffness irrespective of any perturbations in the anchor stiffness during maturation. Our findings offer new insights on biohybrid systems that are essential for their design and an understanding of their emergence.

## RESULTS

### Biohybrid swimmer design

#### Design overview

Our novel swimmer is designed to convert muscle contractions into large amplitude angular displacements of a pair of thin tails to generate high amplitude bending waves and propulsion (Fig. 1). It is also designed to support muscle maturation through mechanical and chemical cues. The swimmer consists of a soft, polydimethylsiloxane (PDMS) scaffold, a ring of skeletal muscle made from C2C12 mouse myoblasts, and a cluster of mouse embryonic stem cell (mESC) derived motor neurons. The muscle tissue functions as the on-board actuator. The scaffold consists of (*1*) a head with a reservoir for neuron spheroids, (*2*) a compliant mechanism with anchors for a muscle ring, and (*3*) two slender tails. A ring of compacted myoblasts is mounted on the anchors. The ring matures with time while it receives mechanical cues from the compliant mechanism and chemical cues from the nearby neurons on the head. Upon actuation, the muscle contracts, pulling the anchors towards each other which results in translation and angular displacement of the base of the tails. The dynamics of the tails generate net time-averaged thrust (〈*F_prop_* 〉) for the swimmer. For high thrust, we need large amplitude muscle contractions which requires a strong, mature muscle. The compliant mechanism is intended to provide an appropriate mechanical stiffness for optimal muscle maturation and yet sufficient compliance to allow for maximal angular displacement of the tails. The wide head of the swimmer increases drag against swimming but also supports a long muscle strip and minimizes the hydrodynamic interactions between the tails by increasing their separation distance. At low Reynolds numbers, the time averaged velocity of the swimmer, denoted by (〈*v*〉, is the time average of the thrust divided by the total drag on the untethered system *γ*, given by 〈*v*〉 = 〈*F_prop_* 〉 / *γ*. We approximate the drag on the head (*γ _head_*) to be that of a sphere at Stoke’s flow with the same projected area in the direction of swimming.

**Fig. 1.**
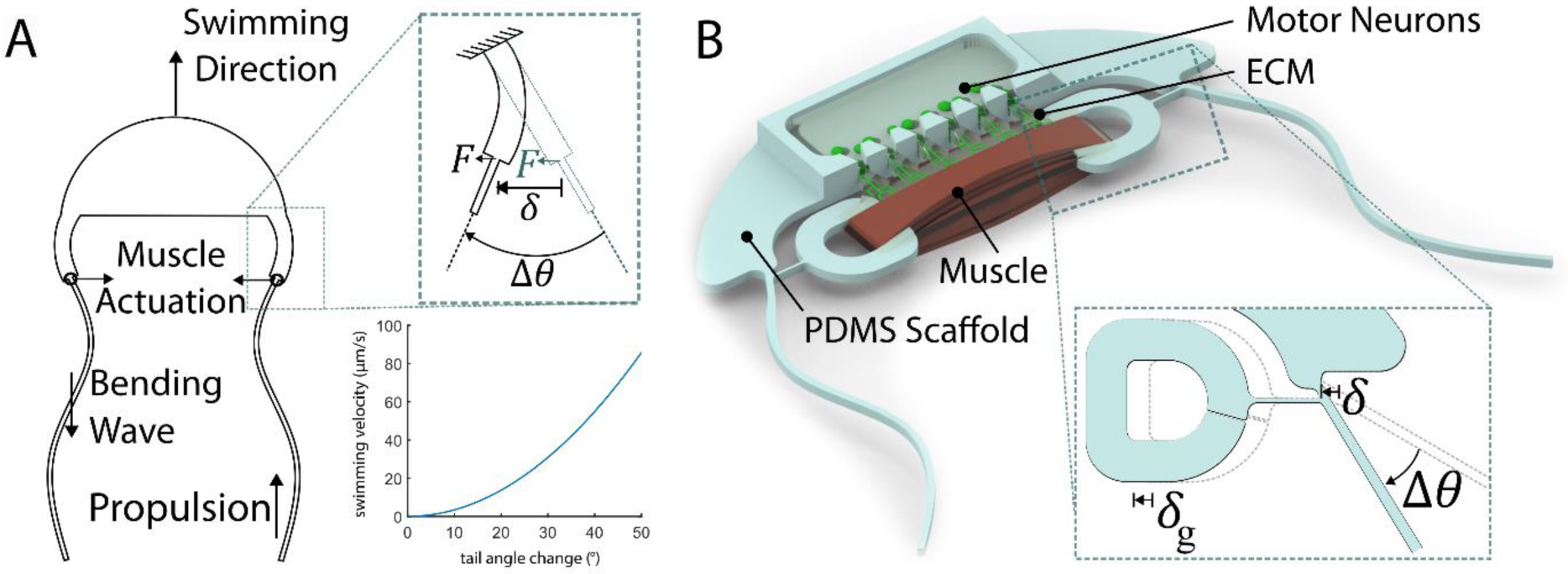
Design for the novel biohybrid swimmer. The amount of curvature in the swimmer tails is exaggerated in these images to convey the idea of a propagating wave. In theory, the optimal speeding speed for a low-*Re* swimmer occurs when the amount of curvature in the tail is approximately one-quarter of a full wave. (**A**) Conceptual design highlighting the role of large angular displacement of the base of the tails for generating large amplitude bending waves and propulsion. The relationship between swimming velocity and tail angle change is derived from numerical simulations of the elastohydrodynamic model of the swimmer. (callout) A conceptual compliant hinge mechanism, whereby the motion of a muscle contracting and relaxing by distance δ creates an angular displacement Δθ at the base of the tail. (**B**) Rendering of the biohybrid implementation of the conceptual design. (callout) The micromolded PDMS realization of the compliant hinge mechanism, whereby the grip is attached to the tail by the thinnest beam which can be fabrication with DRIE and demolded successfully (around 20 µm in our experience).

At low *Re*, thrust generated by a thin filament comes from the asymmetry between the drag per unit length tangent to the filament (*ζ*) and perpendicular to the filament (*ζ* _⊥_), such that *F_prop_* ∝ (*ζ* _⊥_ − *ζ*) (*32*). From resistive force theory we have that in the limit of the length of the filament to the radius of the filament *L* / *r* → ∞, the perpendicular and parallel drag coefficients are related by *ζ* _⊥_ = 2*ζ* (*39*). In our design, where *L* / *r* ≍ 90, we expect *ζ* _⊥_ = 1.6*ζ* (*33*). When the tails are oriented along the direction of swimming, they contribute to the drag of the overall system *γ _tails_* = 2*Lζ*. The overall drag of the system is then *γ* = *γ _head_* + *γ _tails_*.

#### Muscle tissue design

Anchoring a muscle tissue to a synthetic scaffold requires some type of interface. This could be a direct interface where the muscle fibers and collagen adhere to the scaffold or an indirect interface where the muscle wraps around the scaffold. Forming a direct interface requires providing an anchor for the muscle and collagen to adhere to, which has been achieved with PDMS for various cell types through treatment with oxygen plasma followed by incubation in extracellular matrix (ECM) (*40*, *41*). This process of ECM adsorption creates anchorage for the tissue. In our experience, this method is less reliable with muscle tissues, likely because the muscle fibers produce high, localized forces which can overcome the adhesion between the adsorbed ECM and the muscle fibers. In both cases (direct or indirect interface) muscle tissue emerges from a mixture of ECM and myoblasts where the myoblasts collectively compact the matrix around a structure to form a band or a ring (*18*, *23*, *42*, *43*). Muscle maturation is facilitated by spontaneous twitching and tension that are necessary for the formation of cross-striated sarcomeres. Myoblasts compact the gel, approach each other and fuse after contact when multinucleated nascent myotubes emerge. They develop into immature myofibrils with loosely organized actin-myosin structures (*35*, *44*, *45*). Maturation begins after the tissue compacts. In the case of a muscle ring, the tissue is transferred onto the scaffold before it starts to differentiate. In this study, we opted to form the muscle around pillars since this eliminates the need for a complex system of molds which would go around the scaffold. We found that rings with inner diameter greater than 3 mm could be manually transferred onto the swimmer scaffold consistently.

In our scaffold design, the grips are spaced 4 mm apart. For a scaffold thickness of 150 µm, a muscle ring with inner diameter 3 mm will be able to wrap around the grips with only 1.1 mm of slack which gets taken up as the muscle rings continue to compact and mature. Having this slack aids with mounting, whereas too much slack will require more compaction by the tissue before it starts pulling on the grips.

For identifying the appropriate stiffness for the muscle grips to support maturation, we recall a previous study that found optimal contractility in muscles anchored between two pillars with an overall stiffness of 2 µN/µm (*23*). The tissues in this study measured around 500 µm in length with a cross-sectional area around 300 µm^2^. We hypothesize that this optimal stiffness might scale with the geometry of the tissue. To illustrate, let *L*_0_ be the free distance between the muscle grips. And assume that the muscle shrinks by Δ*L* while it matures. Also suppose that the grips have a linear spring constant *k* so that the passive tension in the muscle is *k*Δ*L*. The ratio of the tensile stress *σ* = *k*Δ*L* / *A_c_* to the tensile strain = Δ*L* / *L*_0_ is then *kL*_0_ / *A_c_* where *A*_*c*_ is the cross-sectional area of the tissue. Our hypothesis implies that muscles with given *kL*_0_ / *A_c_* will have similar maturity. Keeping this constant with respect to the previous study gives a stiffness of 14 µN/µm per grip for the current swimmer design.

#### Swimmer tails

The swimmer tails attach to the head of the swimmer via a compliant mechanism which translates the muscle contraction into angular displacement. The two tails cancel out each other’s thrust normal to the direction of swimming. Like the scaffold head, the tails contribute drag to the system, which at low *Re* can be estimated with resistive force theory (*39*). In the limit *Re* → 0, and for an arbitrarily slender tail, there exists a difference between the drag per unit length along the tail *ζ* and perpendicular to the tail *ζ* _⊥_. This difference is referred to as drag asymmetry, and it is necessary for the generation of net propulsion at low *Re* (*32*). Because *ζ*_||_ is less than *ζ* _⊥_, the tails contribute the least drag to the overall system when their resting configuration orients along the direction of swimming. During development, the muscle generates tonic tension, pulling the tails towards the middle of the swimmer. To compensate for this, the tails are initially oriented 60° away from the muscle. For a 4 mm long muscle which shortens by *4*% of its length during electrical stimulation (*12*), the absolute displacement for each grip (anchoring the muscle) would be roughly 80 µm. This displacement of the grips *δ _g_* is converted by the compliant mechanism into linear displacement *δ* and angular displacement Δ*θ* of the base of the tails (Fig. 1B). For fixed actuation amplitudes and tail cross section, the propulsion generated by the tails is fully determined by their sperm number, *Sp*, which varies with length, actuation frequency, and flexural rigidity. For low *Re* swimmers, propulsion is negligible for *Sp* < 1 and is at a maximum around an *Sp* of 2.1 (*32*). The present swimmer is designed for an *Sp* of 1.4 during 5 Hz actuation.

#### Motor neurons contribution to muscle performance

Evidence has been emerging that in vitro muscle matures better in coculture with motor neurons, potentially due to interaction mediated by mutual chemical cues. Neurons extend neurites towards the muscle, some of which may form neuro-muscular junctions. A study of the abdominal muscles of drosophila, which resemble the cross-striated skeletal muscle in mammals, found that tension within nascent myotubes precedes and is required for their development into immature myofibrils. These immature myofibrils exhibit calcium dependent twitching, and this twitching appears necessary for the self-assembly of organized, cross-striated sarcomeres (*35*). Studies have found increased spontaneous twitching in skeletal muscle in the presence of motor neurons. Aydin et al. (2020) developed a platform for the fabrication of C2C12 derived muscle tissues in the presence of mESC derived motor neurons. They observed preferential outgrowth of neurites towards the muscle, higher spontaneous contractility and a 60% higher ratio of cross-striated myofibers to immature myotubes in co-cultured samples. Andersen et al. (2020), developed a system for culturing cortical, spinal, and muscle organoids together. Several times more spontaneous twitching events were observed in the muscle in the presence of the spinal organoid, although the variability was high sample to sample. Transmission electron microscopy images of the multi-organoid samples show highly organized, cross-striated sarcomeres, which is evidence of maturity.

To improve maturity of the swimmer muscle through neuronal signals, we dispense mouse stem cell derived motor neurons on a PDMS holder affixed to the head of the swimmer. The holder also increases the rigidity of the head against bending and buckling out of plane. We fill the holder with mouse embryonic stem cell (mESC) derived motor neuron spheroids.

The holder keeps the motor neurons around 1 mm away from the muscle tissue to avoid any contact between neuron and muscle cell bodies. Studies have shown that spheroids of motor neurons in contact with skeletal muscle tissue form localized, functional neuromuscular junctions (*24*, *46*), resulting in weak, localized muscle activity. In contrast, neurons positioned away from the muscle formed NMJs and exhibited robust tissue-wide contraction (*23*, *47*). In our design, ECM is used to bridge the gap between the motor neuron spheroids and the muscle. Neurites extend through the ECM bridges towards the muscle while the holder prevents the neuron spheroids from slipping onto the muscle during or after placement (Fig. 1B). When ECM is deposited around a muscle tissue, we observe that the muscle tissue pulls the ECM towards itself over several days. In addition, when motor neurons innervate muscle, they build up tension in their axons (*48*). Without mechanical anchorage, this force might pull the neurons into the muscle should innervation occur.

### Implementation and characterization

#### Scaffold fabrication and calibration

The scaffold is molded in a PTFE coated microfabricated silicon mold. The neuron spheroid holder is cast in a separate SLA printed resin mold and is attached to the scaffold with liquid PDMS. For the purpose of analyzing its deformation response, the swimmer scaffold has been sub-divided into four components: the grip, the coupler, the beam, and the head (Fig. 2A). The mechanical stiffness cue experienced by the muscle is measured by pulling the grip of the swimmer scaffold with a tungsten wire with known spring constant (0.43 µm/s, Fig. 2B). The displacement of the head (*x*_1_), bending of the small segment of the tail connecting the head and the coupler (*x*_2_), the stretching of the coupler (*x*_3_), and deformation of the grip itself (*x*_4_) all contribute to the compliance felt by the muscle. Note that the bending of the tail near the base gives rise to the desired angular displacement (Fig. 2C). Initially, a sacrificial support attaches the grips to the head of the swimmer, resulting in an overall stiffness of 1.83 µN/um per grip, behaving linearly up to 400 µN. When the sacrificial support is removed, stiffness drops to 1.75 µN/µm at each grip at small deformations. The stiffness decreases with increasing deformation, dropping to 0.97 µN/µm at 400 µN, i.e., the grip behaves as a softening spring (Fig. 2D). If the grips are released several days after mounting, then the muscle is subjected to a sudden drop in stiffness during maturation followed by further softening of the grips. We show that muscle reaches similar maturity regardless of when the sacrificial support is released, indicating that muscle maturity is invariant to the stiffness history of the anchor. The effect of such change in stiffness during muscle maturation has never been explored before. We discuss this in detail in a following section. The rate at which the tail undergoes angular displacement with respect to the muscle exhibits slight nonlinearity, beginning at around 0.09°/µm when *x*_4_ = 0 and decreasing to 0.05°/µm when *x*_4_ = 400 µm (Fig. 2E). This drop in the angular displacement rate works against the expected swimmer performance, since after maturation muscle already shows a steady grip displacement of 330 ± 29 µm (n = 20, ±SEM). The reason for this appears to be the moment induced by the coupler on the tail hindering its ability to twist with respect to the head.

**Fig. 2.**
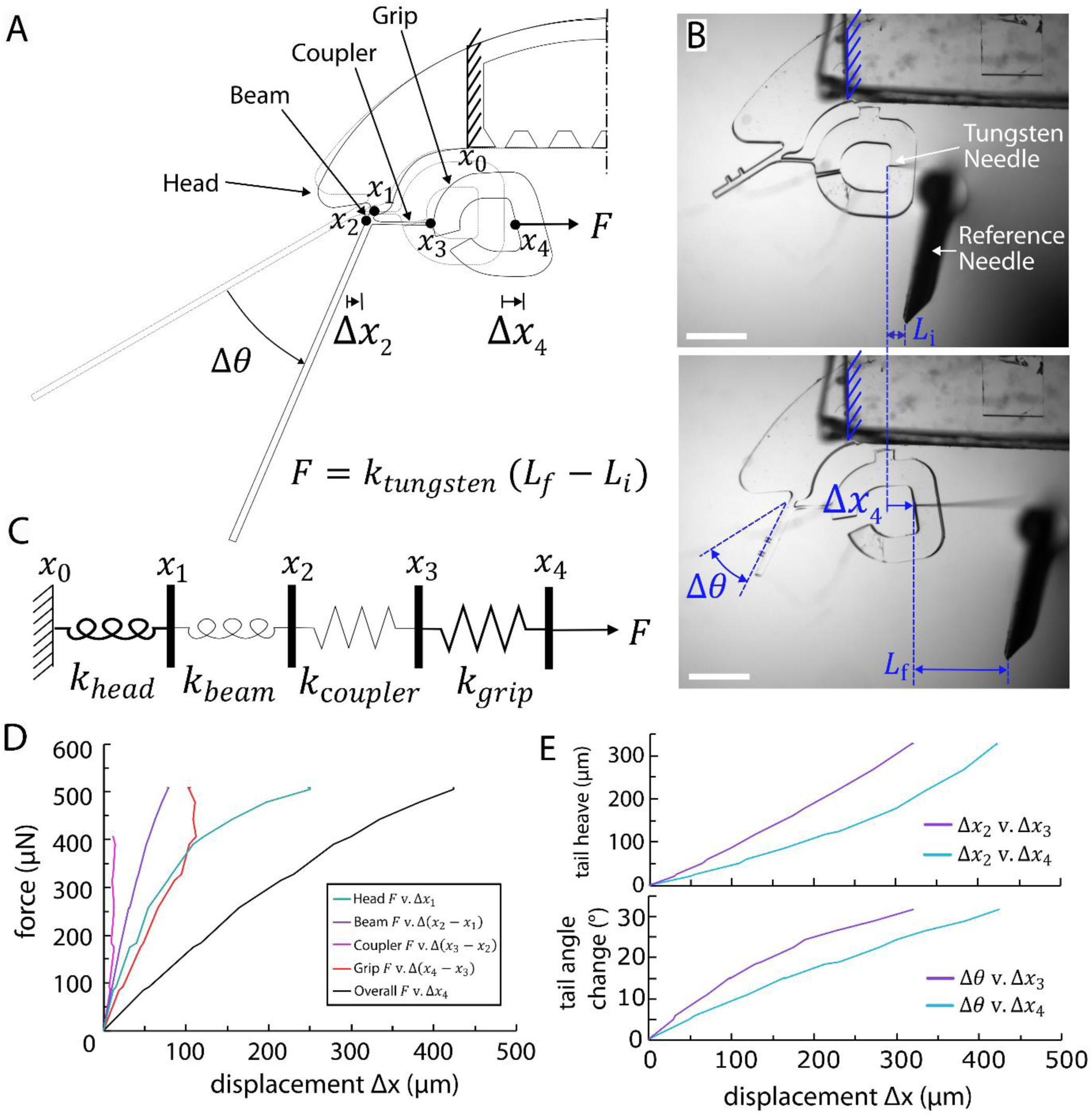
Calibration of the PDMS swimmer scaffold. (**A**) Schematic of the swimmer scaffold decomposing the overall system into four compliant components. (**B**) Diagram of the setup for calibrating the PDMS scaffold with respect to the stiffness of a tungsten needle (0.43 µN/µm) with a hypodermic needle acting as a reference. (**C**) Diagram of the compliant components of the scaffold. The head and the beam contribute to the angular displacement of the tail. (**D**) Force-displacement relationships for the components of the scaffold. (E) Tail angle-displacement relationship which applies to contraction of the muscle.

#### Timeline for biological component assembly

The timeline of the fabrication and assembly of the biological components of the swimmer appears in Fig. 3. The muscle ring fabrication and mounting process (Fig. 3C) are shown in Supplementary Video 1. During mounting, sacrificial supports attach the grips to the head of the swimmer. They offer rigidity against grip displacement and facilitate mounting of the muscle. In order to initiate C2C12 myoblast fusion into nascent myotubes, the concentration of serum in the media is reduced, following the protocol in Olson (1992).

**Fig. 3.**
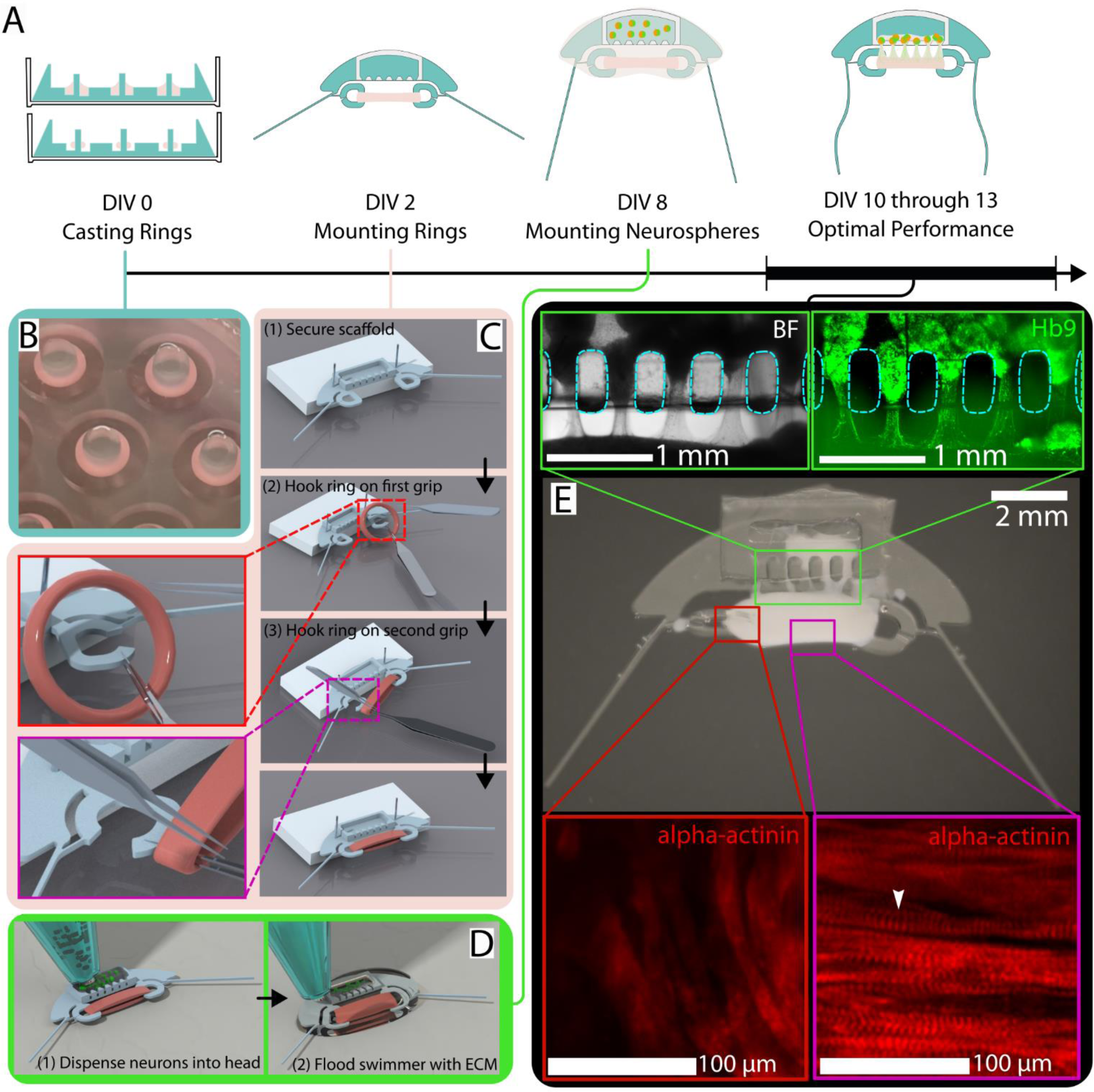
Timeline for the fabrication and assembly of the biological components of the swimmer. (**A**) Overall timeline for the biological components. Some preparation steps occur beforehand, including the expansion of a culture of C2C12 myoblasts and the differentiation of mouse embryonic stem cells into motor neuron spheroids (see Materials and Methods). (**B**) Ring molds used for casting the muscle tissue rings. (**C**) Renderings of the steps involved in mounting the muscle rings on the scaffold. Callouts show tweezer placement necessary to put the muscle into the grips without causing damage. (**D**) Renderings of the steps involved in mounting the motor neuron spheroids in the head of the swimmer. A hydrophobic parafilm substrate keeps the ECM from flowing away. (**E**) Photograph of the swimmer with callouts showing the live imaging of the growth of neurites towards the muscle and confocal images of α-actinin assembled into sarcomeres (white chevron) localized in the middle of the muscle tissue taken after fixing. The neurite GFP signal indicates expression of the Hb9 motor neuron specific promoter, although other types of neurons are present in the neurospheres due to imperfect differentiation efficiency. The photograph of the swimmer was post-processed by darkening the background around the edge of the PDMS scaffold to help the clear rubber body stand out.

#### Muscle maturation is unaffected by the history of the stiffness of the anchor

Although the circumference of the muscle is long enough to wrap around the anchors without deforming the scaffold, in practice, the tissue possesses some rigidity and resists being stretched out. It exerts around 100 µN of force on the anchors following mounting. On day in vitro (DIV) 2, the swimmers are placed in low serum (2% HS) differentiation media. Over the following two weeks, the muscles continue to compact, developing tonic tension against the anchors. In the presence of the sacrificial supports (Fig. 5A) when the compliance of the head, the tail and the coupler are turned off, the muscle is subjected to a stiffness of 1.9 µN/µm per grip (Fig. 4A, 4B). After the supports are cut, the stiffness drops. The amount it drops depends on the force in the muscle. On DIV 3, cutting the supports drops the stiffness to 1.5 µN/µm per grip. On DIV 6 or 9, the stiffness drops to 0.8 µN/µm per grip. We wanted to know if the time history of the stiffness felt by the muscle would have a long-term effect on the rate and extent of the build-up in tension in the muscles. To test this, three groups of swimmers were prepared with muscle rings (no neurons). The sacrificial supports were cut 1, 4, and 7 days after starting differentiation media (DIV 3, 6, 9). The length of the muscle is measured with respect to the points joining the grips and the couplers (*x*_3_ to *x*_3_) (Fig. 4C).

**Fig. 4.**
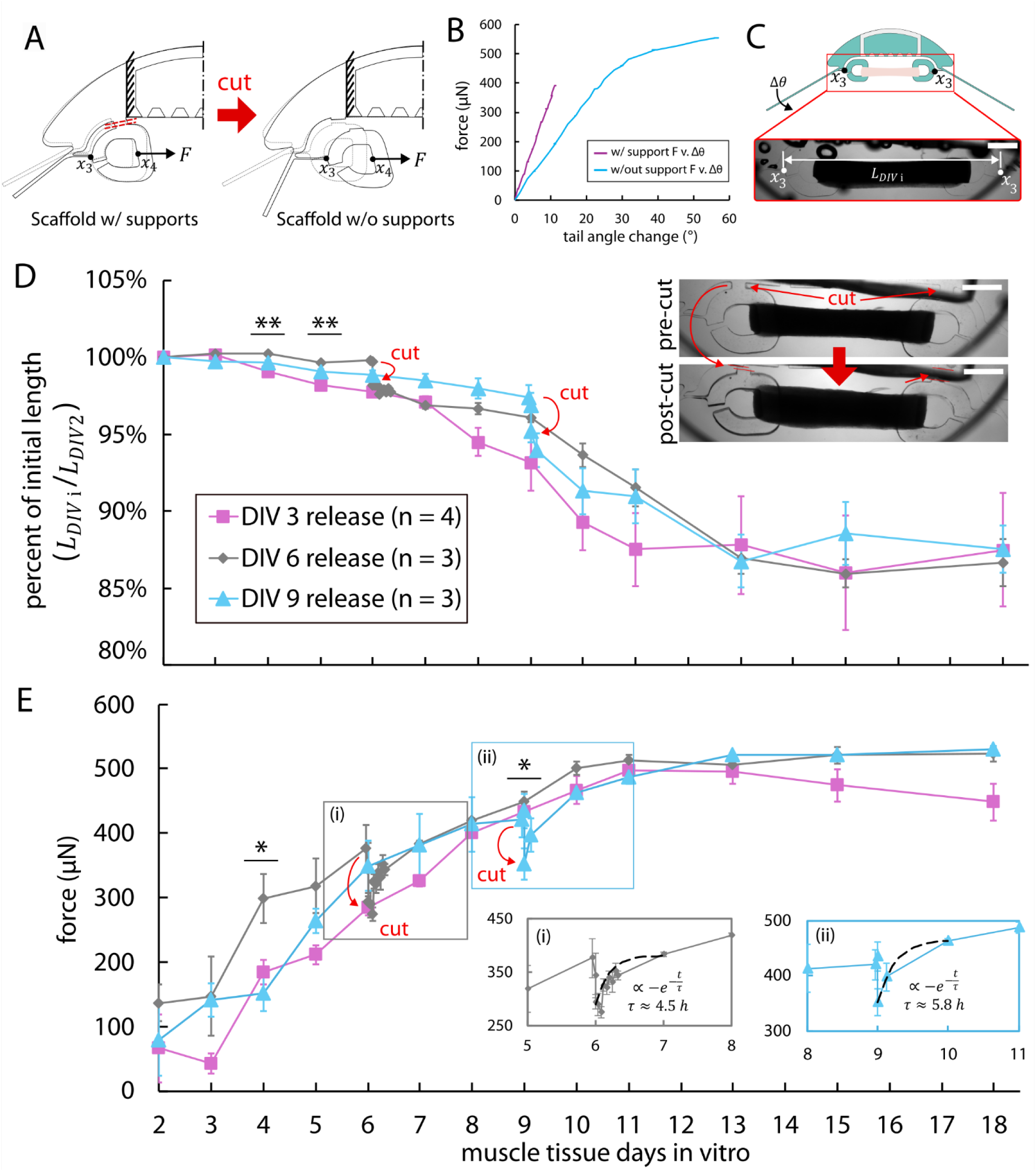
Contraction and tonic force increase of the muscle during development. (A) Schematic showing the change in scaffold geometry following removal of the sacrificial supports. (B) Force-displacement curves showing the stiffness measured from *x*3, the junction between the grip and the coupler before and after cutting the sacrificial supports. (C) Diagram defining how the contraction of the muscle is extracted from bright field images. (D,E) Tracking the variation in the contraction and tonic force of the muscles when their sacrificial supports are removed at DIV 3, 6, and 9. Statistics indicate the results of one-way ANOVA statistical tests comparing the percent of initial lengths of the muscle of the three groups (** = p-value < 0.01, * = p-value < 0.05, n.s. = p-value > 0.05). For DIV 6 and 9, statistics reflect length measurements taken one hour after the removal of the sacrificial supports. By DIV 9, after cutting the sacrificial supports on the last group of samples, large variation among the muscle lengths contributes to the groups being indistinct from one another (p-value > 0.05 by one-way ANOVA). (Ei,ii) insets for change in force following release on DIV 6 and 9. Exponential curve fits give approximate time constants of 4.5 and 5.8 hours for recovery of the force during the first day after release.

**Fig. 5.**
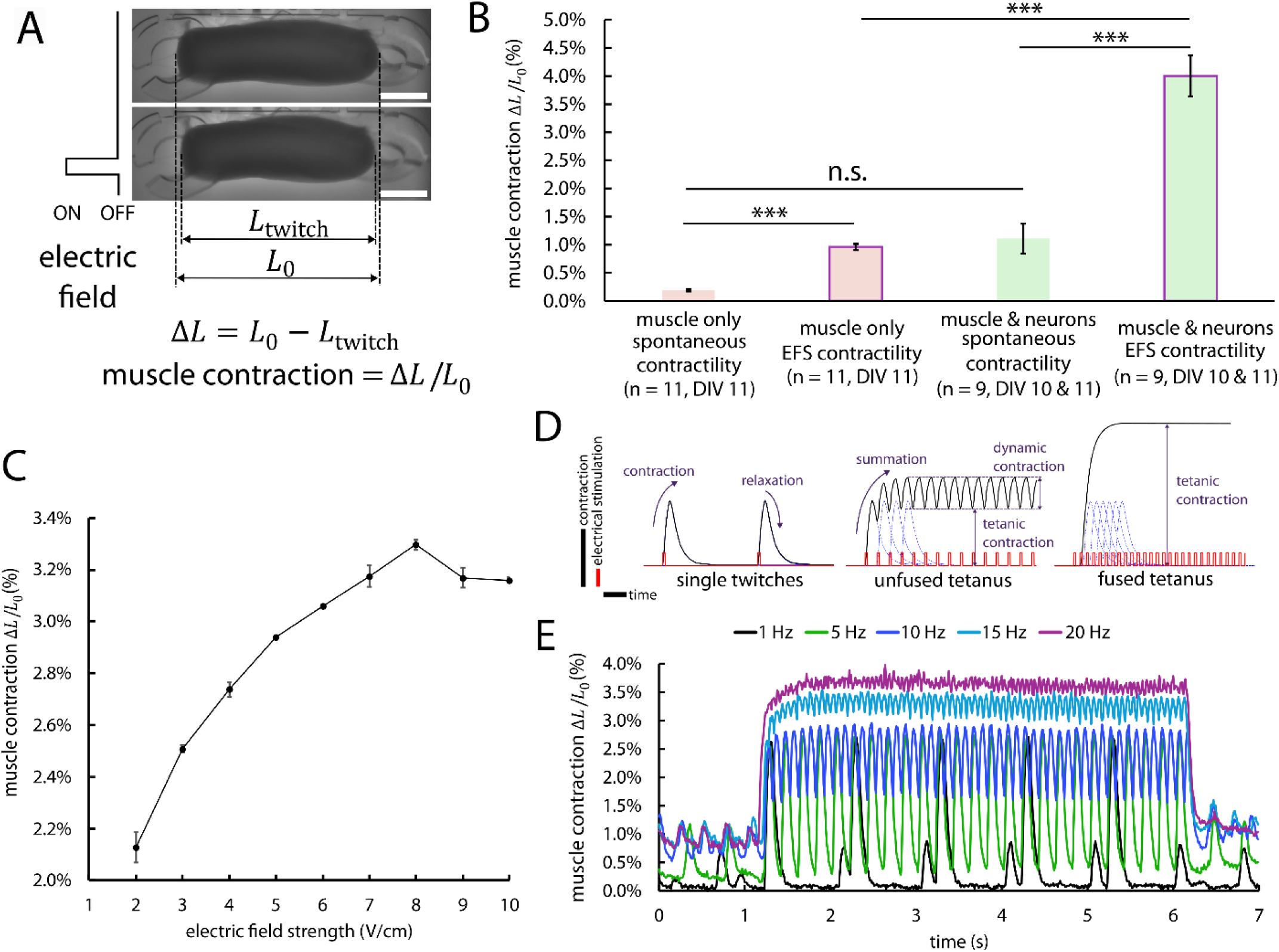
Characterization of the spontaneous and evoked contractility of the muscle tissues. (A) Diagram for how the muscle contractility is measured with respect to the resting length of the muscle. (B) Comparison of the spontaneous and evoked contractility of swimmers with and without the inclusion of motor neurons. Statistics refer to t-tests assuming independent samples (*** = p-value < 0.001, n.s. = p-value > 0.05, error-bars indicate SEM). (C) Variation in contractility with increasing electrical field strength for a typical swimmer. Values refer to the mean of the contraction peaks observed when the muscle is stimulated for 4 pulses at 1 Hz for each voltage (error-bars indicate SEM of the peak values). (D) Diagram of nomenclature used to describe muscle contraction. Muscle contraction follows membrane depolarization and calcium influx. And because the refractory period for the depolarization of muscle is 2-3 ms (*54*), the contractions of a muscle in response to membrane depolarizations can overlap. At low frequency (≤ 5 Hz), a muscle can relax between depolarizations, leading to single isolated twitches. At higher frequencies, the muscle cannot fully relax between depolarizations, leading to a constant shortening referred to as tetanic contraction. The muscle is said to exhibit unfused tetanus. At very high stimulation frequencies (> 20 Hz), the muscle contraction appears smooth and sustained, and the muscle is said to exhibit fused tetanus. (E) Variation in the muscle dynamics with increasing frequency for a typical swimmer. Muscle is subjected to electrical stimulation at each frequency in ascending order, at a field strength of 3V/cm, and with 10 second rest between stimulations. Low amplitude spontaneous twitches (∼1%) can be seen interspersed with the evoked spikes. At higher frequencies, the muscle is unable to fully relax between stimulation frequencies and exhibits spontaneous twitching at increased frequency with comparable maximum contraction amplitude (∼1%). High frequency and high field strength both tend to excite spontaneous twitching, which may indicate the dysregulation of the calcium cycling in the muscle fibers.

When the sacrificial supports are cut, the muscles shorten and their tonic force drops (Fig. 4D,E). The muscles rebuild this tension within a day (Fig. 4Ei,ii). Throughout the development process, the lengths and tonic force of the muscles in the different groups follow a similar course. Before DIV 5, the overall stiffness felt by the groups varies by less than 25%, which may contribute to this similarly. However, even on DIV 8, when the stiffness varies by 90% between the samples with the without the sacrificial supports, the tonic force in all the samples is insignificantly different. Except on DIV 4 and on DIV 9 after cutting the supports in the DIV 9 release group, the tonic force in all the samples is insignificantly different. The different anchor stiffnesses do not appear to affect the tonic force development, and the variation in the time history of the stiffness felt by each of the muscles does not appear to have left a lasting effect on tonic force or length of the muscles. This might be the result of a homeostatic state of tension that the muscle arrives at during maturation, independent of the support stiffness and corresponding force history. On DIV 11, when the muscles would be used for swimming, the average tension in all of the muscles was 498 ± 8.5 µN (n = 10, mean ± SEM).

Starting between DIV 6 and 8, the muscle rings tend to exhibit small amplitude, spontaneous twitching. This indicates that immature myofibrils or matured cross-striated myofibers have developed. To increase the amount of spontaneous twitching, motor neuron spheroids are added on DIV 8. To do this, the swimmers are placed on wax paper, excess media is aspirated, and the motor neuron spheroids are added to the holder on the swimmer’s head (Fig. 3D). Next, the neurons and muscle are covered with ECM. This protects the muscle and neurons from drying out while the ECM polymerizes. When the swimmer is lifted away from the wax paper, some ECM stays attached to the narrow gaps between the muscle and the holder, while excess ECM sticks to the wax paper and tears off. Over the next two days, loose ECM gets pulled into the muscles by the C2C12 cells, and the ECM between the holder and the muscle forms a series of tense bridges. Neurites grow along these bridges towards the muscle (Fig. 3E). By DIV 8, swimmers with neurons tend to exhibit strong, spontaneous twitching on the order of 1.11± 0.27% (n = 9, mean ± SEM) the length of the muscle. This twitching causes the tails of the swimmer to move enough to see with the naked eye (Supplementary Video 2). As a note, the serum concentration is the same for all samples after DIV 2 (2% HS), but the basal media varies between aneural and cocultured samples with DMEM in the aneural samples and a 50:50 mixture of advanced-DMEM/F-12 and Neurobasal Plus media in the cocultured samples.

Concerning the long term stability of the swimmers, it is found that beyond two weeks, the performance of the swimmers declines until they no longer respond to electrical stimulation. One contributing factor to this decline is likely the persistent proliferation of C2C12 inside the muscle tissue, acting to increase its compressive stiffness without adding to the functional muscle fibers. Another factor may be the excretion of cysteine proteases by the C2C12 cells, leading to the breakdown of the tissue. Previous studies have shown that culturing muscle with E-64, a broad-spectrum cathepsin inhibitor, can extend the lifespan of C2C12 tissues to more than 200 days (*49*).

#### Motor neurons increase spontaneous and evoked muscle contraction

By DIV 10, swimmers with and without neurons tend to reach their peak dynamic contractility in response to electrical stimulation. Muscle contractility is measured as a percent change in the muscle’s resting length (Fig. 5A). The amplitude of spontaneous and evoked twitching is compared between swimmers with and without neurons (Fig. 5B). On DIV 10, only 4 of 11 swimmers without neurons exhibit spontaneous twitching, as compared to all 9 of the swimmers with neurons. During electrical stimulation, the swimmers with neurons contract 4.2 times more than the swimmers without neurons on average. Contractility is observed to increase with increasing voltage up to 5 V/cm (Fig. 5C). However, voltages above 3V/cm are sometimes observed to excite prolonged, spontaneous twitching, which hampers our ability to turn off the movement of the swimmer.

Contractility at low frequency is around 2.8% of the muscle length (Fig. 5D). At stimulation frequencies above 4 Hz, the muscle is unable to fully relax before the next pulse, resulting in tonic contraction and decreased actuation amplitude.

We used motor neurons expressing ChR2, a light sensitive ion channel which opens in the presence of blue light (465 nm) and induces the neurons to depolarize (*50*, *51*). At DIV 10, the swimmer samples with neurons were subjected to blue light using the setup from Aydin et al. (2019). In previous studies, stimulation of optogenetic motor neurons has been able to cause global depolarization of an associated muscle tissue (*12*). In those cases, this may have been due to widespread innervation of the muscle by the neurons or due to gap junction (found in immature skeletal muscle) between the myofibers capable to transmitting the wave of depolarization (*52*, *53*). In our samples, there were no significant changes in the spontaneous twitching pattern of the muscles during or after optical stimulation of the neurons, nor in the amplitude of contraction of the muscle. This suggests that any NMJs that might exist are too few in number or too immature to cause the global depolarization of the tissue observed in other studies. It is worth noting that the neurons only contacted a small area on one side of the 0.8 mm^2^ thick muscle tissues. The total volume of neurons measures on the order of 1 mm^3^ as compared to the volume of the muscle which is around 4 mm^3^. We are not able to tell whether the neurites could penetrate into the interior of these muscle tissues.

### Swimming experiments

#### Analyzing swimming velocities

Swimmers exhibiting strong contractility (>2%) to stimulation at 3 V/cm are placed in a dish containing a mixture of coculture media and Percoll (P1644, Sigma Aldritch). The Percoll induces a density gradient, allowing the swimmer, slightly denser than water (1.06 g/mL), to float at a desired height in the media (above the floor and below the air-media interface) due to buoyancy (*55*). Of the twelve swimmers prepared following the timeline in Figure 3, eleven exhibit a response to electrical stimulation, of which six exhibit forward swimming. Swimmer displacement is extracted from videos using an ImageJ template matching plugin (*56*) (Fig. 6A). Points tracked on the body of the swimmer tend to oscillate with the muscle contractions, with the overall path of the swimmer moving along a straight trajectory (Fig 6B). The projection of the swimmer trajectory in the direction of travel is used to extract the velocity of the swimmer before and during electrical stimulation (Fig. 6C). Contraction of the muscle is calculated by tracking a point on the grip with respect to the head of the swimmer and then adding the displacement of the grip due to the tonic contraction of the muscle. At rest, the swimmers are observed to drift at speeds around 1-2 µm/s.

**Fig. 6.**
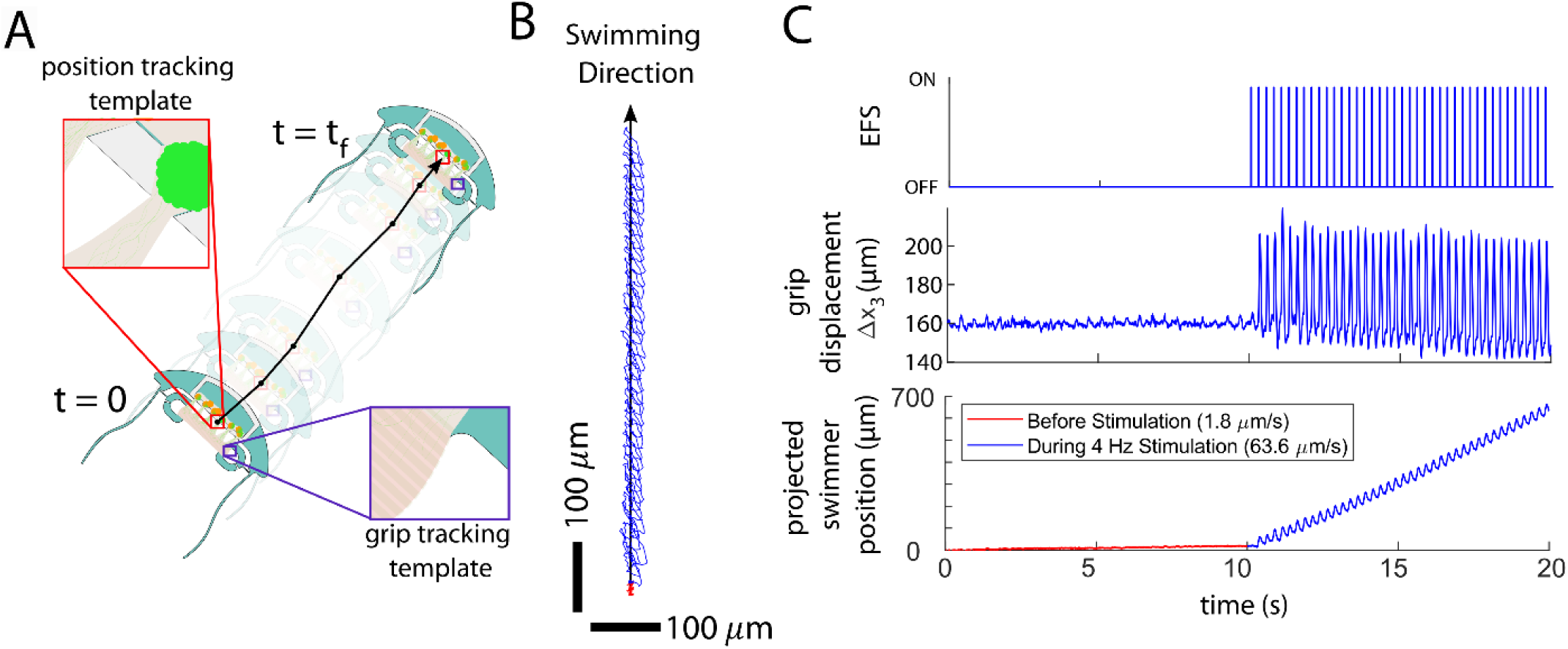
Swimming position tracking and analysis. (A) Diagram of the template tracking method employed to extract the swimmers position and muscle contraction waveforms. (B) 2D projection of the path of a point on the head of the swimmer before (red) and during (blue) 4Hz electrical stimulation. (C) The corresponding stimulation, grip displacement, and position of the swimmer during the 4 Hz electrical stimulation. The position is taken by projecting the swimmer’s path onto the direction of travel. Before stimulation, the swimmer drifts with a small velocity. This is attributed to convective currents in the dish.

This is attributed to convective currents in the media. In the example (Fig. 6B,C), the swimmer moves with a time averaged velocity of 63.6 µm/s, which corresponds to a *Re* of 0.33 (Supplementary Video 3). During each muscle twitch, the base of the tails are subject to an average angular displacement of 4.8 ± 0.04° with a peak angular displacement rate of 176 ±1.6° / *s* mean ± SEM). (n = 39, At higher frequencies, the muscle undergoes tetanus as it is unable to relax between stimulation pulses. The onset of tetanus brings the tails together, giving the swimmer a sudden high velocity kick. Similarly, when the electrical stimulation ends, the release of tetanus allows the tails to move apart, which causes a high velocity kick backwards. These transients are excluded from the calculation of time averaged swimming velocity by only fitting the swimmers position over time from one second after the start of stimulation till the end of stimulation.

#### Persistent motion of the swimmer

We varied actuation frequency (0.5 to 20 Hz) and measured swimmer dynamics (Fig. 7A-D) (Supplementary Video 4). At frequencies 5 Hz and below, where the muscle experiences minimal tetanic contraction, the swimmer appears to reach a steady time-average velocity within a few cycles. And once the stimulation ends, the swimmer appears to continue moving forward while decelerating (Fig. 7E*i*). At frequencies ≥ 8 Hz, the initial onset of unfused tetanus causes the swimmer to move forward rapidly at a speed of 178 ±16 µm/s (n = 30, mean ± SEM). Over 1-2 seconds, the swimmer appears to decelerate towards a steady, non-zero time averaged velocity, which we attribute to the small amplitude dynamic contractions of the muscle (Fig. 7E*ii*). Once stimulation stops, the muscle relaxes back to its baseline causing the swimmer to move backwards at a speed of 159 ±13 µm/s (n = 30, mean ± SEM) before decelerating (Fig.7E*iii*). Across all samples during 0.5 Hz stimulation, the muscle length during contractions changes at a peak rate of 1.54 ± 0.10 mm/s and relaxes slower, with a peak rate of 1.31± 0.14 mm/s (n = 18, mean ± SEM). This may account for some of the difference between the forward and backward velocities of the swimmer due to tetanus.

**Fig. 7.**
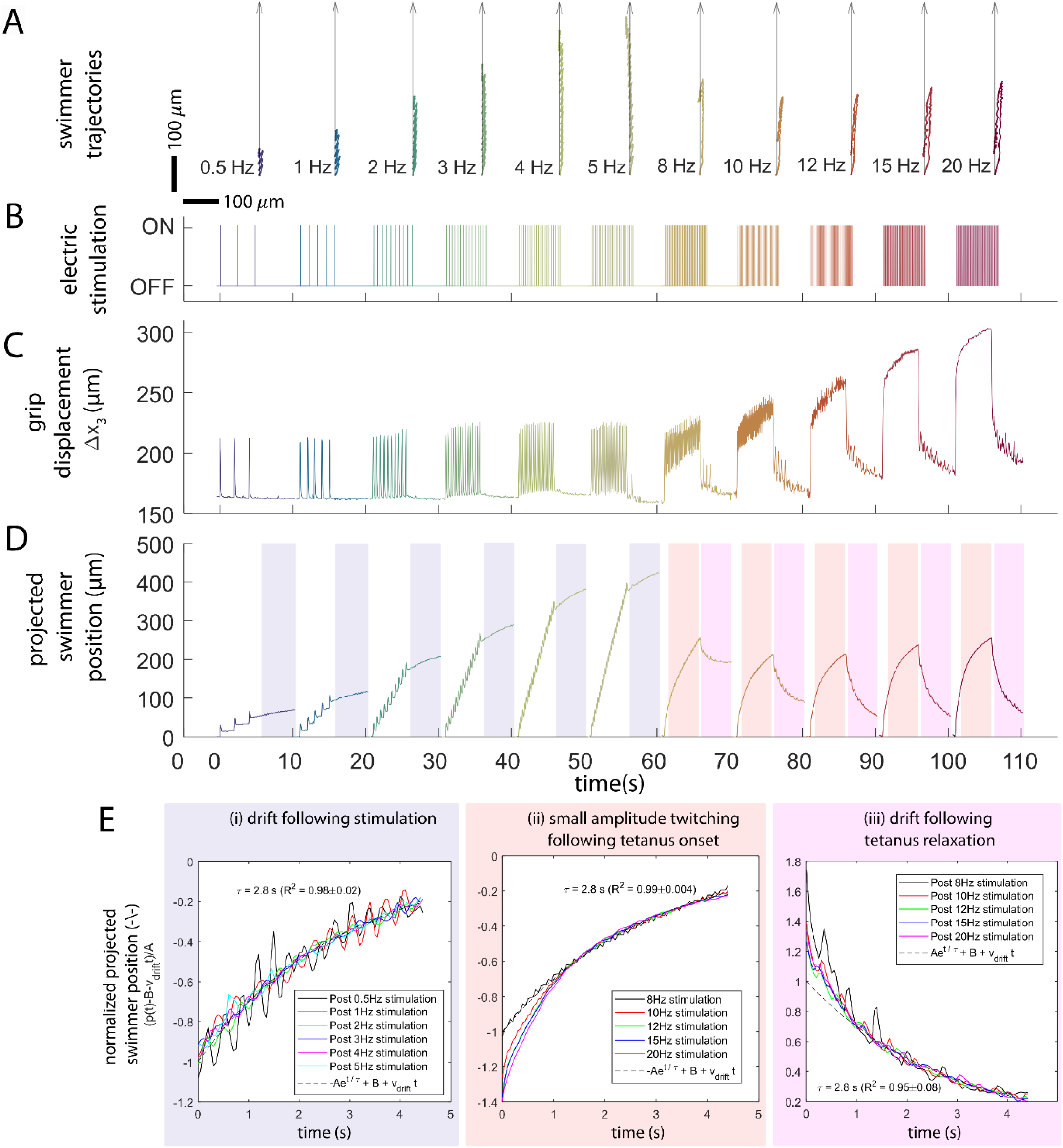
Swimming waveforms for a high performing swimmer. (**A**) x-y coordinate plots for the swimming trajectory of the swimmer at multiple frequencies. (**B**) Electrical stimulation waveforms (3V/cm, 10 ms pulse width). (**C**) Muscle contraction measured from motion of *x*_3_, the junction between the grip and the coupler. At frequencies above 5 Hz, the muscle experiences significant tetanic contraction. (**D**) swimmer position calculated by projecting the trajectory in the direction of swimming. At frequencies ≥ 8 Hz, the onset and relaxation of tetanus gives the position waveform a distinct sawtooth shape. (**E**) Nondimensionalization of the swimmer dynamics with time constant τ = 2.8s for each of the cases considered. Background colors refer to time windows in (D). During these time windows, we hypothesize that the dominant force on the swimmer is drag. They include (i) when the muscle is almost motionless, so that the only force exerted on the swimmer is drag, (ii) when the initial propulsion from the onset of tetanus is large compared to the propulsion from subsequent dynamic contractions, and (iii) when the propulsion from tetanus relaxation is large compared to the subsequent, slower muscle relaxation.

We investigated the source of the swimmer’s deceleration with a model of ballistic motion which only accounts for the drag force. Suppose the swimmer has mass *m*, and experiences drag proportional to its velocity with a constant drag coefficient γ. And suppose that the swimmer has some initial velocity *v*_0_ and position *x*_0_ at time *t*_0_. Let *x*(*t*) be the position of the swimmer over time in the lab reference where the fluid is drifting with some small velocity *v_drift_*. From the force balance, *mx* = −*γ* (*x* − *v_drift_*). Solving the initial value problem gives *x(t) = v_0_m/γe^t−t_0_/m/γ^+v_drift_(t − t_0_) + x_0_*. The characteristic time constant τ for the deceleration is mass divided drag *m* / *γ*. We fit experimentally measured *x*(*t*) of the swimmer to the above equation for the following 3 cases: (i) soon (1 s) after low frequency (0.5 – 5 Hz) stimulation is stopped, (ii) during high frequency (8 – 20 Hz) stimulation when muscle reaches tetanus state with low amplitude contraction, and (iii) soon (1 s) after high frequency stimulation is turned off. The delay of 1 s is to avoid any transient dynamics soon after applying or removing stimulation. All the data in Fig. 7 comes from a single swimmer with a wet mass of approximately 27 mg. Based on the speed of the swimmer at the start of the experiment, the drift velocity *v_drift_* is approximately 1.76 µm/s. Each curve fit for each case yields a slightly different value for the time constant τ, with an average value of 2.8 seconds. Using τ = 2.8s, we repeat the fitting to calculate values for *v*_0_ and *x*_0_. Using these values for *v*_0_ and *x*_0_, each of the curves can be nondimensionalized to isolate the first-order dynamics (Fig. 7E).

The velocities seen for this example swimmer range from 10 µm/s to 80 µm/s, which corresponds with *Re* ∼ *O*(10^-1^). The value of *Re* can be thought of in terms of a ratio of the contribution of inertial forces and viscous forces in the hydrodynamics of a body. At these intermediate values (*Re* ∼ 1), we would expect both the inertial and viscous forces to be present in the propulsion generating mechanism of the swimmer. Whereas viscous forces contribute to propulsion through drag asymmetry of the tails, inertial forces would contribute by imparting momentum to the surrounding fluid. This regime of *Re* is not considered for flagellate swimmers (*12*, *32*).

For a wet mass of 27 mg, the average time constant of 2.8 s corresponds to a linear drag coefficient of 9.6 mg/s. This is lower than the predicted drag for the two tails, *γ _tails_* = 21.6 mg/s. This is significantly lower than our full prediction of the drag of the swimmer that includes the head drag, γ = 42.6 mg/s (for a projected forward cross-sectional area of 3 mm^2^). In the following section, a low *Re* model of the swimmer dynamics is provided with the tail boundary conditions from our experiments. The model is used to predict the swimming velocities. This calculation assumes that the swimmer comes to rest immediately once the propulsion force *F_prop_* from the tails is removed, such that the velocity *v*(*t*) = *F_prop_* / *γ*.

#### Comparison between experimental swimming and low Re simulations

We plot the velocities of our six swimmers as a function of tail angle amplitude. Swimming velocities increase with increasing tail angle across aggregated data from all six swimmers (Fig. 8A). In the case of frequencies which induced tetanic contraction, the tail angle change corresponds with the dynamic contraction and excludes the tetanic contraction (Fig 5D). In some cases, the swimmer was observed to move backwards during stimulation. Across the frequencies tested, peak swimming velocity was observed at 5 Hz. In order to assess agreement with our low *Re* elasto-hydrodynamic model (*33*), swimmer velocities were calculated with drag coefficients γ = 9.6 mg/s and 42.6 mg/s, and experimental muscle contraction waveforms for frequencies in the range of 0.5 to 5 Hz (Fig. 7C) and for Δ*θ* ∈(0°,8°] (see Methods and Materials). Neither drag coefficient yields velocity predictions above 40 µm/s. When considering a single swimmer performance at different velocities, we find model predictions are several times slower than experimental values (Fig. 8B). This suggests more thrust is being produced than predicted by low *Re* dynamics. This increased thrust and speed is likely contributed by inertial effects. The evidence for this comes from the deceleration of the swimmer from any arbitrary initial velocity with a single time constant in the absence of any actuation or external force.

**Fig. 8.**
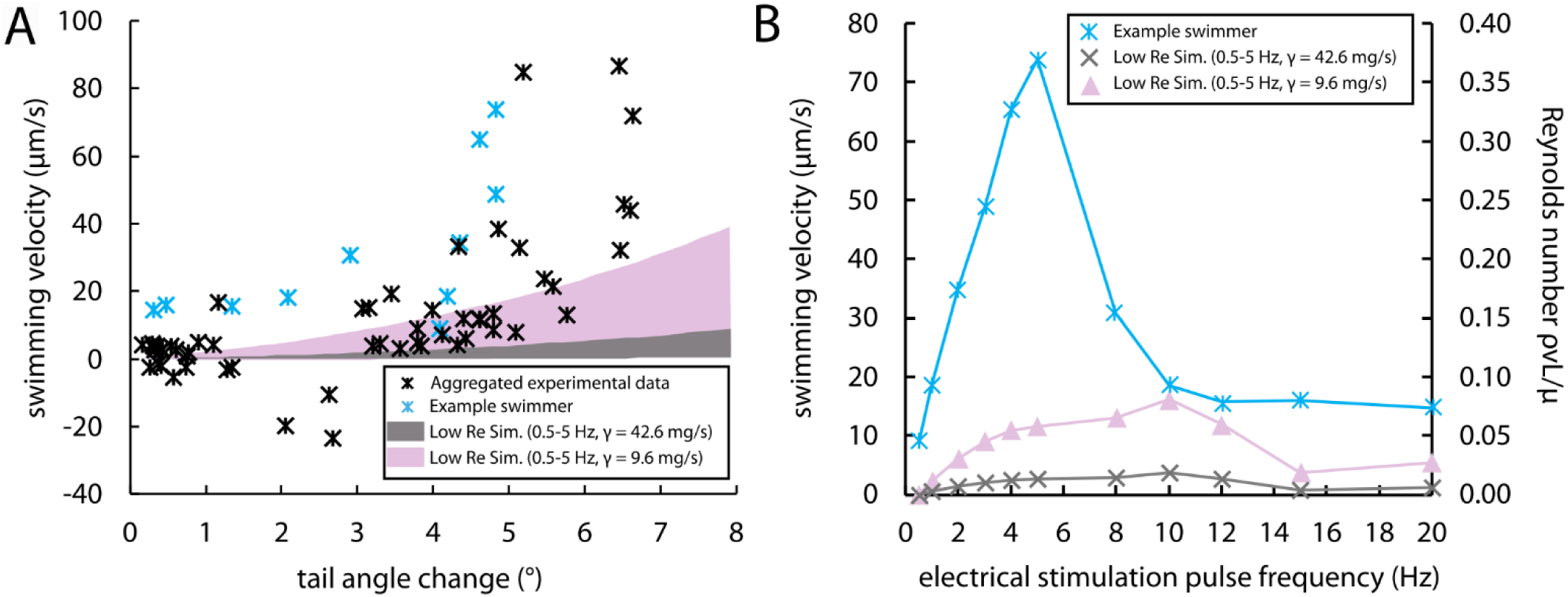
Comparison between experimental swimmer velocities and predictions from our low *Re* model. (**A**) Aggregated swimming velocities and tail angle changes across all samples (n = 6) against regions corresponding to model predictions generated with the indicated frequencies, tail angle changes, and drag coefficients. Tail angle changes correspond with the dynamic contraction (Fig. 5D) of the input waveform, excluding tetanus contraction. Simulations are run using experimental muscle contractions waveforms as the model inputs. Inputs are scaled to achieve the associated tail angle change. (**B**) Comparison of experimental and predicted swimmer velocities at multiple stimulation frequencies for a single swimmer. The predicted swimming velocities are generated using the experimental muscle contraction for this swimmer as the model input.

## DISCUSSION

In this study, we fabricate and test a muscle-powered biohybrid swimmer designed to swim at low *Re* using the non-reciprocal motion of a pair of tails to generate net-thrust. Our design focuses on supporting the maturation of a muscle through mechanical and chemical cues to provide a large input displacement to a compliant mechanism which subjects the base of the tails to angular displacement. When electrically stimulated, swimmers reach velocities two orders of magnitude higher than previous biohybrid flagellate swimmer designs. When stimulation stops, the swimmers are observed to drift forward under their own inertia. Low *Re* elasto-hydrodynamic simulations made to mimic our experiments predict speed around four times slower than are observed. Taken together, our results suggest that inertial forces, expected at intermediate *Re* ∼ *O*(*1*), are playing a role in the dynamics generating thrust by our swimmer.

### Adaptability of the muscle to attachment stiffness

The tonic tension in the developing muscles grows at a similar pace irrespective of when the sacrificial supports are removed, with the only significant difference between the three groups being observed on DIVs 4 and 9 following the removal of the sacrificial support in the last group (Fig. 4E). We observed the tonic tension build-up in three groups of muscles over a 2-week period The muscle builds up tonic tension against a constant or softening stiffness. When the stiffness drops due to the removal of the sacrificial support, the muscle quickly contracts to reach the same level of tonic tension it had before the drop. We interpret this as the muscle adapting to its changing environment to reach a homeostatic tension. This tension appears to increase up to DIV 15 before leveling off. A comparable result was observed in Powell et al. (2002) with tissue-engineered human skeletal muscle. On the days between DIV 9 and 14, researchers applied a constant stretch to the muscle, increasing it by 5% every two days. And despite the perturbation, the tonic tension in the muscle stayed in the range of 400 to 600 µN. This is similar to the tonic tension found in our muscles, and by coincidence, our muscle rings and their tissues have similar cross-sectional areas of roughly 1 mm^2^. In their case, applying a constant stretch has the effect of increasing the tonic force in the muscle, and the muscle is able to relax and re-establish its homeostatic tension. In our case, the muscles are able to rebuild their tonic tension after drops in tension and changes in stiffness.

The drop itself is only 25-90% depending on the day of release, and in all cases, the stiffness less than 14% of the hypothesized optimal stiffness of 14 µN/um. We suspect that for a range of “soft” stiffnesses, the increase in tonic force of the muscle will follow a similar time-course which is robust to changes in the stiffness to other values within this range.

Robustness to changes in attachment stiffness within a range would allow for increased flexibility in the design of biohybrid machines. With the ability to adjust to perturbations in stiffness, muscle-powered machine can be fabricated with sacrificial supports that allow for the system to gradually reach a final configuration. Already, supporting structures are a common element in biohybrid system (*8*, *18*, *58*). Our results do not indicate how different stiffnesses will impact other aspects of the muscle aside from tonic force. A comparison between muscle development on soft and stiff pillars has been attempted before in the context of training in vitro tissue constructs with electrical stimulation (*59*). It was found that the benefits of training with electrical stimulation depended on the anchor stiffness. On a soft pillar, the training leads to increased contractility, but on a stiff pillar, training over multiple days had no improvement over stimulating the muscle for 1 hour on the last day of the test. So, although tonic forces may be comparable for different stiffnesses, the performance of the muscle and its response to other cues may vary with stiffness. In addition, it has been found that muscles cocultured with motor neurons exhibit hypertrophy from chronic electrical stimulation while muscle tissues by themselves do not (*60*). We would therefore suspect that the performance of skeletal muscle and its response to training will depend on anchor stiffness and whether it is cocultured with motor neurons. Such an investigation would benefit from granular tunability of the spring stiffness and traditional measures of muscle maturity including mRNA markers (*60*), sarcomere numbers, and ECM composition (*61*).

### Effects of motor neurons on muscle performance

Muscles grown in coculture with motor neurons demonstrate several-fold higher contractility compared to those grown by themselves (Fig. 5B). Although neurites are observed to grow to the muscle (Fig. 3E), the improvement to contractility occurs within two days of adding the neurons. This may be too soon for the formation of mature NMJs, which takes three to five days in vitro (*12*, *62*, *63*). Considering the asymmetry between the volume of neurons and the volume of the muscle (1:4), as well as restrictions on where the collagen bridge connects the neurons to the muscle in our swimmer design, we suspect that the improvement in muscle contractility is due to chemical cues from crosstalk between the cell types. In our samples, the neurites were able to sense and grow towards the muscles, which may have been due to their proximity and the presence of a physical bridge of ECM between them. Although functional improvement in muscle due to coculture with neurons has been observed many times, the coculture parameters necessary for this improvement are unclear, and could be studied with this platform by adjusting the proximity and intermediary between the muscle and neurons.

### Inertia in swimmer propulsion

We observed persistent motion of the swimmer following stimulation, following the onset of tetanus, and following the release of tetanus. This is evidence of momentum which is not expected at ultra-low Reynolds numbers. Experimental velocities are 3-4 folds higher compared to predictions based on Stoke’s flow (Fig. 8). This is not surprising since *Re* ∼ 10^-1^ found in our experiments is much higher than that usually considered in the Stoke’s flow regime, for example, *Re* ∼ 10^-3^ for sperm dynamics (*64–66*). If inertial forces are contributing to the propulsion, this would mean that the tails of the swimmer are imparting momentum to the fluid. Because the density of the swimmer is almost identical to the density of the fluid, evidence of inertial drift of the swimmer (Fig. 7E) suggests inertial drift of the fluid.

These intermediate Reynolds numbers have been examined theoretically in the literature for some special cases (*67–70*). Their authors looked at the speed of a swimmer with respect to its flapping Reynolds number *Re_f_* = *ρ_body_ AωL* / *μ*, calculated from the frequency *ω* and amplitude *A* of actuation, and the body density *ρ_body_*. Theoretical studies predict swimming speed proportional to *Re*^α^_f_ for dense swimmers (body density *ρ_body_* >> fluid density *ρ _fluid_*,*α* = 1) and swimmers near asymmetric boundaries (*α* ∈{1/ 2,1}). In both cases, the introduction of inertia to the propulsion is continuous with increasing *Re*. The applicable case to our swimmer is one with swimmer density similar to the surrounding fluid (*ρ_body_ ρ _fluid_*), which has not been investigated theoretically for the general case (*69*). However, an analysis was performed for the specific case of a dimer swimmer by linearizing the Navier-Stoke’s equations around *Re* ∼ 1 (*71*) A few biological swimmers have a similar size scale to that of our swimmer, including newborn tadpoles, however they operate at a higher *Re*, on the order of 10^2^ to 10^4^ (*72*, *73*). A species of sea slug called *Clione antarctica* exhibits both non-reciprocal low *Re* and reciprocal high *Re* locomotion. Studies looking at the transition between the two swimming modes have found that inertial propulsion would arise discontinuously when the amplitude and frequency of the flapping of the slugs “wings” exceeded a threshold (*74*). They attributed it to a transition through a hydrodynamic instability. They computed this threshold with respect to the unsteady Reynolds number *Re_ω_* = *ρ ωL*^2^ / *μ*, which is calculated based on the frequency of flapping *ω* and the length scale of the body *L*. It is a measure of how quickly the sea slug flaps its wings. They found that the threshold lies between *Re_ω_* ∈[5, 20], which may be specific to their organism. A more appropriate parameter for our swimmer might be the velocity of the tails during stimulation, which is around 15.5 mm/s at the tip and depends mostly on the muscle contraction speed and less on the stimulation frequency because muscle twitches are self-similar at all frequencies up to the onset of tetanus.

Although our reasons for increasing the input angular displacement to the tails come from low *Re* theory, there are benefits to intermediate *Re* swimming (Fig. 8A). Newborn tadpoles are able to reach velocities in the mm/s to cm/s range by undulating their bodies. With further improvements in our ability to generate large amplitude angular displacements using muscle tissues, it may be possible to design a system which achieves these speeds while remaining at a mm-length scale.

In summary, we demonstrate a novel flagellate-like biohybrid swimmer capable of reaching velocities two orders of magnitude greater than previous flagellar swimmer designs of similar size. The velocities also exceed predictions based on Stokesian flow (low *Re* ∼ 10^-6^). This is attributed to inertial forces contributing to thrust in our swimmers at *Re* ∼ 10^-1^. Higher thrust was achieved by the novel design of the swimmer scaffold which transduces muscle contraction into large angular displacement at the base of the tails. Time average swimming velocity increases sharply with change in the tail angle during muscle contraction. We found that C2C12-derived muscle adjusts its tonic tension towards a homeostatic value, irrespective to the history of stiffness cues it experienced during maturation. This is a critical finding from the study that deserves further investigation to reveal how changing stiffness effects the response of tissue-engineered muscle to other supportive cues, such as electrical, chemical, and mechanical stimulation. Future iterations of the swimmer design may turn towards optimization algorithms involving Finite Element analysis to maximize tail angle change with respect to muscle contractions.

## MATERIALS AND METHODS

### Swimmer scaffold fabrication and calibration

The swimmer scaffold is molded from polydimethylsiloxane (PDMS, Dow Corning Sylgard 184, 10:1 base to crosslinker ratio) at 60°C for two days using a silicon wafer mold for the head and tails (〈100〉 orientation, 500 µm thickness, undoped) and a 3D printed resin mold for the neuron holder. The wafer is patterns with SPR220 and etched to a depth of 155 µm using deep reactive ion etching (DRIE) in an STS Pegasus. It is then coated with PTFE which acts as a release agent that helps with the demolding of PDMS. The resin mold is printed from Clear V4 resin in a Form3 SLA 3D printer with a minimum feature size of around 200 µm. The two components are manually assembled with liquid PDMS and cured at 60°C for two days. The assembled swimmer scaffold is autoclaved in distilled water to drive out monomers and then autoclaved again with a dry cycle to sterilize. The force-displacement relationship at the grips of the scaffold is calibrated in water with respect to a tungsten needle of known length and diameter (Living Systems Instrumentation, Young’s modulus 411 GPa, 0.43 µN/µm stiffness). During calibration, the scaffold is adhered to a dish with double sided tape at the edge of where the neuron holder sits.

### Skeletal muscle ring fabrication and mounting

Rings of C2C12 mouse myoblasts are fabricated and matured following standard protocols (Z. Li et al., 2019; Mancini et al., 2011). Briefly, P4 C2C12 mouse myoblasts are thawed from cryogenic storage and resuspended in growth media (Gibco’s DMEM, 1X GultaMAX, 1X Penicillin/Streptomycin, 10% FBS) and plated at a density of 20k cells/cm^2^ in a T75 treated culture flask. The growth media is changed every other day, and the cells are passaged once reaching 50-70% confluency.

Annulus shaped ring molds with inner diameter 3 mm are fabricated from PDMS using from a SLA printed resin negatives. To prevent cell adhesion, the ring molds are incubated in a 1% Pluronic solution (Sigma-Aldrich Pluronic F-127, reconstituted from powder and filtered through 0.22 µm pores) at 20°C overnight. Afterwards, the solution is removed, and the ring molds are rinsed 3 times with PBS.

A C2C12-ECM mixture is prepared containing 2.5M cells/mL, 2 mg/mL Type I rat tail collagen, and 2 mg/mL Matrigel. The collagen is neutralized with a 7% sodium bicarbonate solution and 10X MEM. Rings are cast from 70µL of the mixture and cured at 37°C for 45 minutes. The mold is then inundated in growth media. The time when the muscle rings are cast is used to demarcate the start of the count of days in vitro (DIV). The rings are left to compact until DIV 2. The compaction ratio is around 10:1, resulting in rings with cross section 1 mm by 0.5 mm and an inner diameter of 3 mm.

On DIV 2, the rings are mounted on the biobot scaffold as shown in Figure 2. Briefly, the biobot scaffold is fixed to a block of PDMS using insect pins, and places under a stereoscope. The scaffold is inundated with growth media, and the ring is added to the dish. Using two pairs of tweezers, the grips of the scaffold are opened, and the ring is placed inside. Then, the media is changed in all dishes to reduce the chance of infection. The rings are maintained in low serum differentiation media until DIV 18 or until motor neurons are added (Gibco’s DMEM, 1X GultaMAX, 1X Penicillin/Streptomycin, 2% HS, 1X ITS). The muscles tend to start responding to electrical stimulation on DIV 6 (monophasic, 10 ms pulsewidth, 3V/cm field strength).

The sacrificial supports connecting the grips to the head of the swimmer are removed using surgical scissors under a stereomicroscope. The scissors cutting into the PDMS causes the grips to twist.

The muscle rings tend to survive this step because they can flex along with the grips. Attempts are making tissues with finger-like grips, like those found in (Emon et al., 2021), fail because the adhesion of ECM to PDMS is insufficient to tolerate this torsion. Once the supports are cut, tonic tension in the muscle rings tends to cause one of the grips to twist 90°.

### Preparation and seeding of the motor neuron spheroids

The procedure for preparation of motor neuron spheroids from optogenetic mouse embryonic stem cells line ChR2H134R-HBG3 Hb9-GFP is described elsewhere (Aydin et al., 2019; Uzel et al., 2016b). The procedure for adding these spheroids to the swimmer is shown in Figure 2. The swimmers are removed from their mounting pedestals and placed on a parafilm sheet with a 100 µL droplet of media. Spheroids are mixed with the same reconstituted mixture of collagen and Matrigel as the muscle rings. The droplet of media is entirely removed from around the swimmer, and the spheroid-ECM mixture is injected into the neuron holder on the head of the swimmer. An additional 20 uL of ECM is dispensed around the head of the swimmer and the muscle to protect it from dehydration and to form a continuous bridge of ECM from the muscle to the neurospheres. The ECM is cured at 37°C and 100% humidity for 30 minutes. Next, the swimmers are pinned back onto their mounting pedestals and inundated with coculture media (47% v/v Gibco’s Neurobasal Plus Medium, 47% v/v Gibco’s Advanced DMEM/F-12, 2% v/v FBS, 1X GlutaMAX, 1X Penicillin/Streptomycin, 1X Gibco’s B-27 Plus Supplement, 1X ITS, 0.1 Beta-mercaptoethanol).

Loose ECM dislodges, but the ECM surrounding the neurospheres and which has infiltrated itself in the narrow space between the muscle and the head of the swimmer tends to remain intact, forming a bridge. This ECM bridge frequently survives reinundation when the sacrificial supports are still intact. The media is refreshed every other day.

### Electric field stimulation and recordings

Typically, for voltages exceeding 1.5V, electrolysis will cause bubbles to form at the electrodes and the pH of the media to shift. However, at sufficiently short pulse-widths, bubbles do not appear to form, and the pH remains stable. In this study we use a maximum pulse-width of 10 ms. Aside from keeping the swimmer healthy, this is important because the formation of bubbles can induce flow in the media which can cause the swimmer to drift. The strength of stimulation is controlled by fixing the voltage across a pair of electrodes normalized by the distance between the electrodes. During stimulation, electrical current passes through the media, and the component which passes through the muscle initiates contraction. The actual voltage drop across the muscle can depend on many factors including the thickness of the tissue and the orientation of the muscle fibers relative to the electrodes.

The performance of the muscle is recorded (Olympus IX81, 2X lens, brightfield, 100fps) subject to electrical field stimulation (monophasic, 10 ms pulsewidth) delivered by a pair of graphite plates (20 mm by 20 mm by 3 mm) positioned on either side of the swimmer. To evaluate swimming performance, a swimmer is placed in a 100×15 mm polystyrene dish (Thermo Scientific, BioLite) containing 30 mL of coculture media and 10 mL of Percoll (Sigma-Aldritch). Using tweezers to break the surface tension, the swimmer is submerged. It is seen to float in the Percoll gradient around 3 mm above the bottom of the dish. The graphite electrodes are placed 4 cm apart to provide clearance for the swimmer. Recordings are taken at 20 or 100 fps, depending on the length of the experiment due to RAM limitations. Additional recordings are taken with a Canon Rebel Ti camera (29.97 fps), which allows for tracking the swimmer over a larger field of view.

### Image processing and analysis

Tracking of the head of the swimmer is performed using the template matching plugin in ImageJ. Swimming displacement is quantified by projecting the motion of the swimmer onto its time-averaged swimming trajectory. For a given test, the trajectory may vary with time to account for rigid body rotation of the swimmer which can result from asymmetric thrust generation by the tails. Once the overall motion of the swimmer has been subtracted, the displacements of the grips are tracked from a point on the grip near the coupler.

### Low Re elasto-hydrodynamic model simulations

The full details of the elasto-hydrodynamic model simulated in this paper appear in Aydin et al., (2024), and the relevant MATLAB codes are available (https://github.com/wdrennan/Biohybrid-Swimmer-Digital-Assets). A second order accurate, implicit finite difference method is used for simulating the filament dynamics. The simulation takes the heave (*x*_2_) and twist (Δ*θ*) of the tails as boundary conditions. These are calculated based on the grip movement at *x*_3_ for the test shown in Fig. 7 using the calibrations shown in Fig. 2E. When simulating the swimming speed for a wider range of angles in Fig. 8A, the *x*_3_ (*t*) waveform for a given frequency is scaled so that the amplitude corresponds with the appropriate twist angle change using these same calibrations. The simulations are run using the experimentally derived drag coefficient (γ = 9.6 mg/s) as well as the predicted drag coefficient (γ = 42.6 mg/s). As a note, in the low *Re* model, the thrust depends on the drag asymmetry of the tails, *ζ* _⊥_ − *ζ*. The experimentally derived drag coefficient (γ = 9.6 mg/s) is smaller than that calculated for the tails (γ = 21.6 mg/s). This experimental derived drag coefficient contradicts our calculated value of *ζ* _⊥_ and *ζ* _||_. In the low *Re* model, we use the predicted values of *ζ* _⊥_ and *ζ* _||_ to compute the thrust, even when this constitutes a contradiction with the experimental drag coefficient. We assume a PDMS stiffness of 1.9 MPa and use the dimensions of our swimmers tail cross-section of 100 µm wide and 155 µm deep (*13*).

### Immunostaining of the muscle rings

For immunofluorescence imaging, samples were fixed in 4% paraformaldehyde (PFA) in phosphate-buffered saline (PBS) overnight at 4°C. The samples were then permeabilized using 0.2% Triton X-100 in PBS and blocked with a solution of 3% bovine serum albumin (BSA) and 5% normal goat serum (NGS) in PBS. Following this, the samples were incubated overnight at 4°C with a rabbit anti alpha-actinin primary antibody (1:250, Abcam #ab68167). After washing the samples five times with PBS (5 minutes each wash), they were incubated with the goat anti rabbit Alexa fluor 568 secondary antibody overnight at 4°C. The next day, samples were washed three times with PBS and incubated with 4′,6-diamidino-2-phenylindole (DAPI) (1:1000, Invitrogen) for 10 minutes, followed by another three washes in PBS. Images were acquired using a Zeiss LSM 700 confocal microscope equipped with an EC Plan-Neofluar 20X/0.5 N.A. objective (Carl Zeiss AG).

### Statistical methods

A series of 3-independent sample, weighted, one-way ANOVA tests were used to compare the force and length of the aneural swimmer in each of the three sample populations on each day (Fig. 4D,E). On days where the sacrificial supports were cut, the force and length values measured after cutting were used. P-values are used to reject the null hypothesis that all the swimmer samples come from the sample population. A series of Students two-tailed t-tests assuming independent samples were used to compare the spontaneous and evoked contractility of aneural and cocultured swimmer samples (Fig. 5B). The behavior of a swimmer during inertial drift was characterized by least-square curve fitting with the given exponential function with affine offset (Fig. 7E). The value for *v_drift_* is taken by fitting a linear function to the projected swimmer position over time before stimulation began. R^2^-values are presented as means ± SEM for the given groupings.

## Supplementary Materials

Supplementary Video 1 – Seeding and mounting C2C12 muscle rings (*3*)

Supplementary Video 2 – The cocultured swimmer samples twitching spontaneously in their petri dishes

Supplementary Video 3 – 4Hz electrical stimulation test of the swimmer sample shown in Fig. 6

Supplementary Video 4 – Frequency sweep test of the swimmer sample shown in Fig. 7 and 8

## Author contributions

Conceptualization: OA, MTAS, ZL, WCD

Data curation: WCD, AB, YK, AC, MW, DD

Formal analysis: WCD, AB, YK

Funding acquisition: MTAS

Investigation: WCD, AB, YK, AC, MW, DD, OA, ZL

Methodology: WCD, BE, MTAS, OA, ZL

Project administration: MTAS, WCD, ZL

Resources: MTAS, OA, BE

Software: OA, WCD, AB, YK

Supervision: MTAS, OA, ZL

Validation: WCD

Visualization: WCD, AB, OA

Writing – original draft: WCD, MTAS, MSHJ

Writing – review & editing: WCD, MTAS, OA, MSHJ

## Data and materials availability

Data used for the figures and analysis appearing in this paper will be made available through the Illinois Digital Environment for Access to Learning and Scholarship (IDEALS) repository (http://www.ideals.illinois.edu/) and is available to reviewers at the time of the submission of this document. Raw *.tif images will be made available through the Illinois Data Bank at the time of publication (http://databank.illinois.edu/). Backups of all digital assets are maintained on hard drives stored in the PI’s lab. Other digital assets including MATLAB codes, CAD drawings, and vector drawings used for the figures are made available on Github (https://github.com/wdrennan/Biohybrid-Swimmer-Digital-Assets).

